# Characterization of morpho-functional traits in mesophotic corals reveals optimized light capture and photosynthesis

**DOI:** 10.1101/2021.09.29.462347

**Authors:** Netanel Kramer, Jiaao Guan, Shaochen Chen, Daniel Wangpraseurt, Yossi Loya

## Abstract

The morphology and skeleton architecture of photosynthetic corals modulates the light capture and functioning of the coral-algal symbiosis on shallow-water corals. Since corals can thrive on mesophotic reefs under extreme light-limited conditions, we hypothesized that microskeletal coral features optimize light capture under low-light environments. Using micro-computed tomography scanning, we conducted a comprehensive three-dimensional (3D) assessment of small-scale skeleton morphology of the depth-generalist coral *Stylophora pistillata* collected from shallow (5 m) and mesophotic (45 m) depths. We detected a high phenotypic diversity between depths, resulting in two distinct morphotypes, with calyx diameter, theca height, and corallite marginal spacing contributing to most of the variation between depths. To determine whether such depth-specific morphotypes affect coral light capture and photosynthesis on the corallite-scale, we developed 3D simulations of light propagation based on photosynthesis-irradiance parameters. We found that corals associated with shallow morphotypes dissipated excess light through self-shading microskeletal features; while mesophotic morphotypes facilitated enhanced light absorption and photosynthesis under low-light conditions. We conclude that the mesophotic coral architecture provides a greater ability to trap solar energy and efficiently exploit the limited light conditions, and suggest that morphological modifications play a key role in the photoadaptation response to low-light.

## Introduction

Biogenic calcification in corals plays a vital role in facilitating reef biodiversity and complexity (Graham and Nash, 2013). Coral calcification comprises the secretion of calcium carbonate crystals in the form of aragonite (Drake et al., 2020), producing a great diversity of geometrical structures and fulfilling the multifunctional purposes necessary to maintain reef health (Zawada et al., 2019). For example, the structural complexity of reef-building corals, on both the reef-scale (m-km) and the coral colony scale (cm-m), provides a broad diversity of habitats for reef-associated organisms. Specifically, small and cryptic fishes, which constitute the main proportion of the coral-reef fauna, rely on the corals’ high structural heterogeneity for their survival (Munday and Jones, 1998; Pereira and Munday, 2016; Wehrberger and Herler, 2014). In addition to genotypic variations, light conditions and water movement are important factors controlling coral geometrical growth (Bruno and Edmunds, 1997; Doszpot et al., 2019; Ow and Todd, 2010; Soto et al., 2018). For some coral species, growth under different environmental conditions can result in changes in their skeletal structure, a phenomenon referred to as “morphological plasticity” (Todd 2008). This phenomenon is believed to be beneficial in enabling such coral species to occupy a wider array of abiotic conditions than those with fixed morphologies (Bruno and Edmunds, 1997; Willis, 1985), and is thus thought to promote the ability of corals to withstand rapid environmental change (Doszpot et al., 2019; Grottoli et al., 2014; Smith et al., 2007).

In particular, it has long been suggested that phenotypic plasticity in corals is advantageous for maximizing light interception and use across a broad range of depths and/or light regimes (Anthony and Hoegh-Guldberg, 2003; Barnes, 1973). Indeed, the relative abundance of different coral morphotypes can often reflect the environmental conditions in which they reside (Chappell, 1980; Doszpot et al., 2019; Dubé et al., 2017; Kramer et al., 2020; Paz-García et al., 2015). For example, the preponderance of plating colonies in mesophotic coral ecosystems (MCEs; characterized predominantly by blue light and 1-20% of surface photosynthetically active radiation (PAR); Laverick et al., 2020), has been attributed to the extremely low light conditions in their surrounding habitat, resulting in their beneficial growth strategy for maximizing incoming light surface area (Kramer et al., 2020). Corals that are exclusively found in either shallow or mesophotic depths are commonly termed “depth-specialists” (Bongaerts et al., 2010). Such corals exhibit permanent morphological modifications acquired through genetic change (i.e., adaptation) that may have evolved to suit local conditions that significantly differ from those of their ancestral origin conditions (Sherman et al., 2019). In contrast, coral species that occupy a broad depth range are termed “depth-generalists”, and are found overlapping between the shallow and the upper mesophotic zones (Bongaerts et al., 2010; Kahng et al., 2014). In essence, a depth-generalist coral species can inhabit light regimes that vary by up to two orders of magnitude (Tamir et al., 2019).

Analogous to patterns in terrestrial plants, variation in light quantity and quality can drive both physiologically and morphologically based strategies for efficient light utilization in corals (Anthony and Hoegh-Guldberg, 2003). In plants, apart from the well-known physiological modifications (e.g. greater quantities of chlorophyll-*a* pigments), leaves in shaded environments are generally thinner and larger as compared to light-adapted leaves (Bragg and Westoby, 2002; Lichtenthaler et al., 2007). Furthermore, the same features can also appear in leaves subjected to the blue spectrum of light (Sæbø et al., 1995). Similarly, depth-generalist corals inhabiting mesophotic environments often exhibit structural modifications that are hypothesized to aid in the utilization of light capture (Einbinder et al., 2009), thereby enhancing photosynthetic performance and optimizing colony growth under limited optical conditions. For example, thinner skeletons and an increased coral tissue surface area to volume ratio is considered energetically more efficient for the capture of incident light when its availability is low (Anthony et al., 2005; Kahng et al., 2020). Thus, modular photosynthetic corals can regulate their internal light regime by varying the extent of self-shading surface on the colony scale towards a photosynthetic optimum (Anthony et al., 2005; Ow and Todd, 2010; Paz-García et al., 2015; Wangpraseurt et al., 2014).

Although understanding the mechanisms that optimize light capture by corals has been the focus of many studies, as far back as the early 1980s (Dubinsky et al., 1984; Dustan, 1982; Falkowski and Dubinsky, 1981), the functional significance of morphology at mesophotic depths has not been thoroughly explored, hindering a comprehensive understanding of the various species’ photoadaptative capabilities. Previous work on photoadaptation at mesophotic depths has been mainly focused on physiological and biochemical alterations (reviewed in Kahng et al., 2019), while most of our understanding of the interaction of coral architecture with light is primarily derived from the whole-colony growth form (Anthony et al., 2005; Einbinder et al., 2009; Hoogenboom et al., 2008; Willis, 1985). Research focusing on the extent to which measurable small-scale morphological traits can be informative regarding the light-harvesting mechanisms employed by scleractinian corals remains insufficient, particularly for corals inhabiting the mesophotic environments.

Recently, 3D-imaging analyses obtained via advanced technologies such as micro-computed tomography (µCT) and laser scanning, have enabled accurate and detailed information on the coral skeleton structure at both the macro- and microscale levels (House et al., 2018; Zawada et al., 2019). Using high-resolution µCT scanning, we sought to determine the role that morpho-functional traits play in light-harvesting. To this end, we assessed the variations in the small-scale skeletal structure of the common depth-generalist coral *Stylophora pistillata* between contrasting light regimes, from the shallow (5 m) and the upper mesophotic (45-50 m) depths in the northern Gulf of Eilat/Aqaba (GoE/A). Based on our morphometric measurements, we conducted 3D light simulations integrating known physiological and optical properties in order to examine the effect of the coral architecture on its photosynthetic performance. Our findings have revealed unique structural intraspecific changes in corals between depths; and we discuss the functional significance of these traits in effectively capturing and dispersing light in their ambient environments. These findings provide a novel understanding of how small-scale morphology-based mechanisms facilitate optimized light-harvesting in MCEs.

## Results

### Skeletal morphometrics

Overall, *S. pistillata* colonies exhibited distinct morphotypes between shallow and mesophotic origins, as determined by PERMANOVA (*p* < 0.001; Fig. 2-4). The first two axes of the PCoA captured 82.6% of the total observed variation in the morphological space between shallow and mesophotic colonies. The first axis explained 71.6% of the variance (Fig. 4) and was most correlated with TH, CD, and CSM (contributing 16.1%, 14.2%, and 13.5%, respectively). Similarly, SIMPER analysis identified that most of the differences in small-scale skeleton architecture were attributed to these same traits, which accounted for over a third of the morphological variation observed between depths. Furthermore, while Pearson’s correlation scores were highest and positive between CD, TH, and SPL, they were negatively correlated with CSM (*p* < 0.01; Fig. S2). Excluding CH and CSC, all morphometric characters significantly differed between shallow and mesophotic specimens (MEPA, *p* < 0.01; Fig. 2). In general, most of the shallow morphological traits exhibited larger sizes compared to their mesophotic counterparts; albeit, with higher variability among the shallow colonies, as seen in the morphospace (Fig. 2, 4). For example, CD was on average ∼60% larger in shallow samples, ranging from a diameter of 0.848 to 1.191 mm compared to 0.533 to 0.719 mm in mesophotic samples (Fig. 2a). In contrast, CSM was greater in mesophotic specimens compared to in shallow ones, exhibiting 58% more spaced corallites (Fig. 2i).

**Figure 1.**
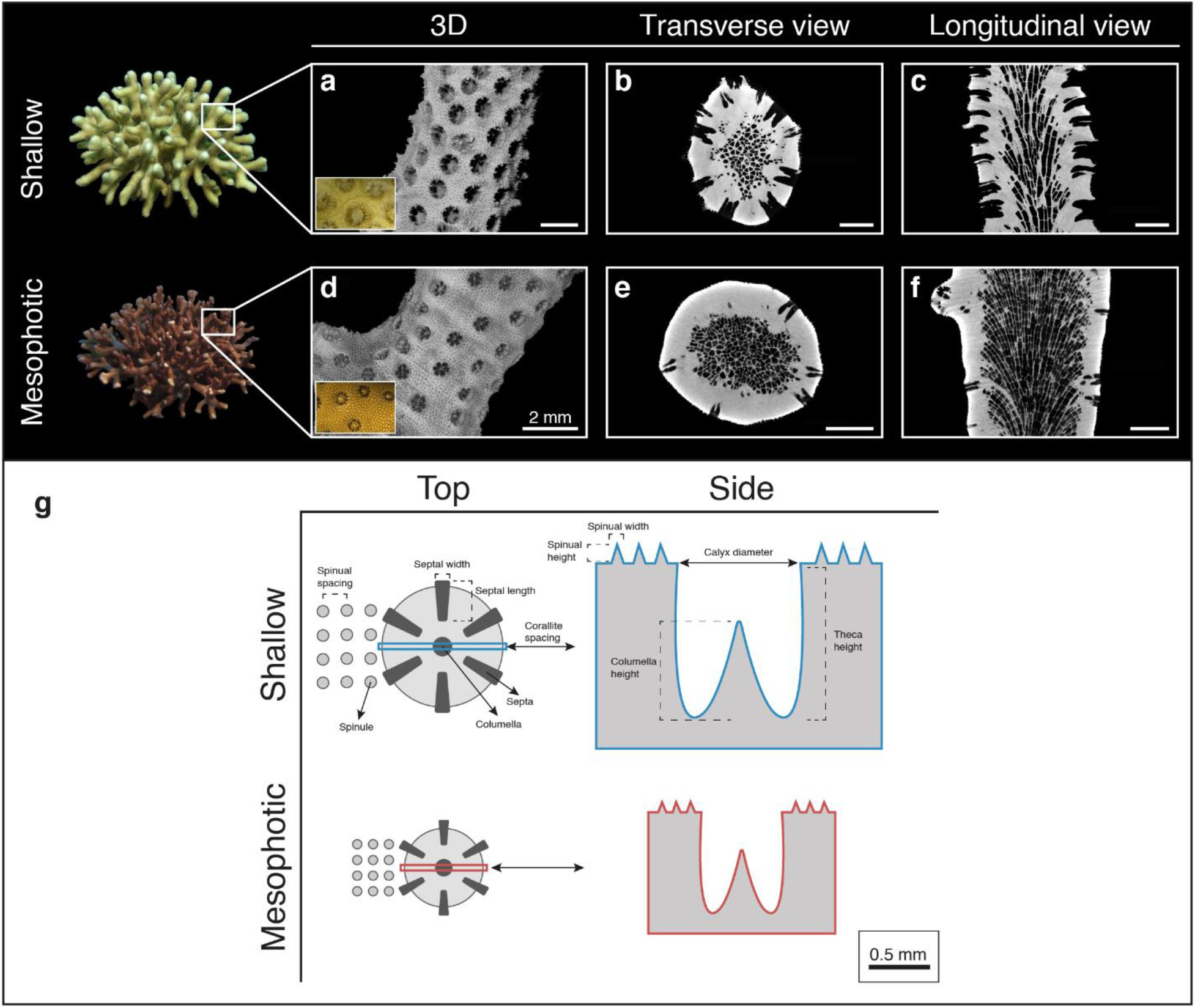
Morphotypes of **(a-c)** shallow and **(d-f)** mesophotic *S. pistillata*. Examples of μCT X-ray scans showing: **(a+d)** 3D reconstructions of the skeletons (inset photos show surface covered with live tissue) and sections of **(b+e)** transverse and **(c+f)** longitudinal scan slices. Scale bars are 2 mm. **(g)** Two-dimensional schematic representation of the top-down and side view of a corallite and its surrounding coenosarc between the studied shallow and mesophotic corals. Key skeletal structural elements are noted and scaled based on mean values.

**Figure 2.**
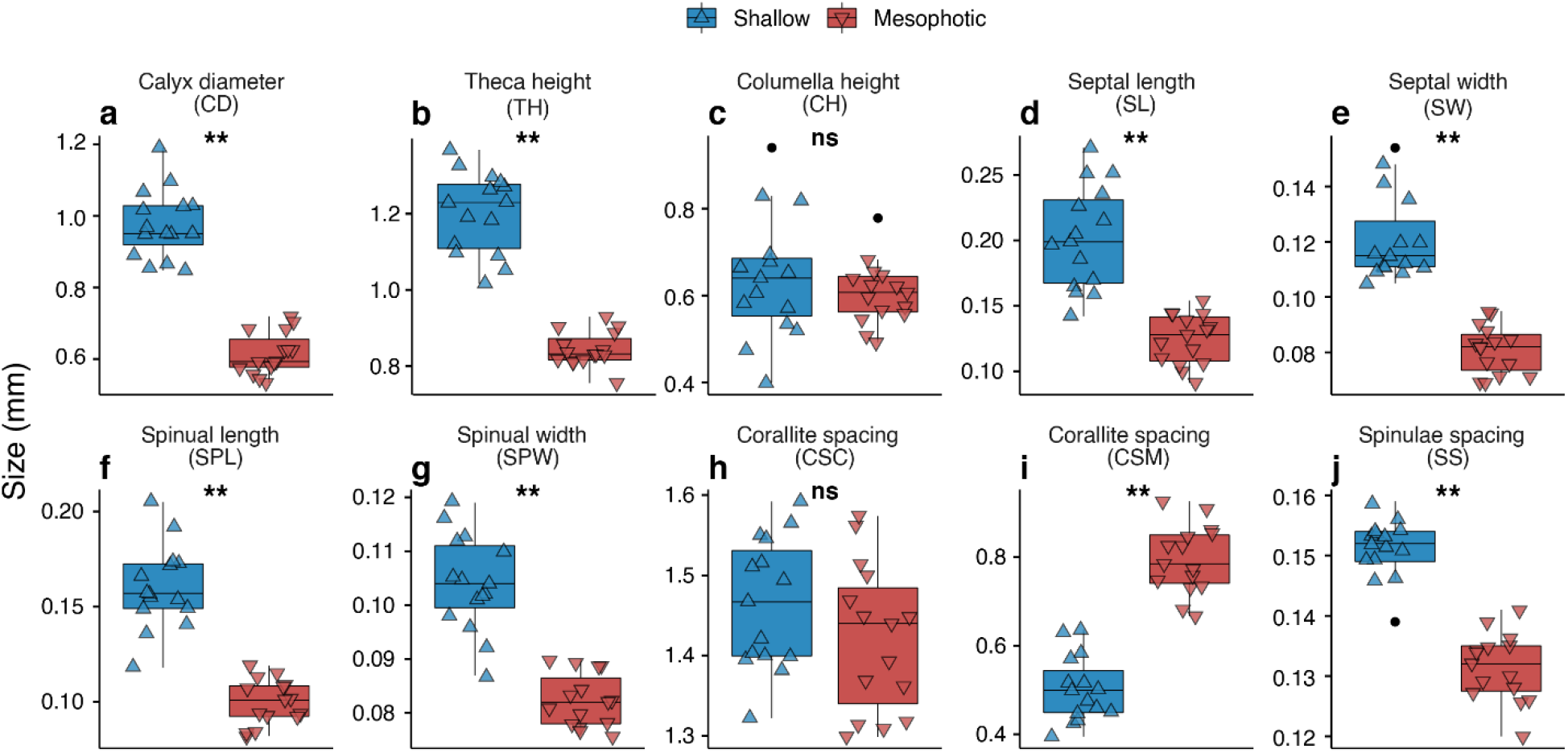
**(a-j)** Box plots showing the mean size variation of morphometric traits between shallow (*blue*; triangle point up) and mesophotic (*red*; triangle point down) *S. pistillata* colonies (*n* = 30). Horizontal lines depict the median, box height depicts the interquartile range, whiskers depict **±**1.5x interquartile range, and dots represent outliers. Asterisk denotes significance (*p* < 0.01).

**Figure 3.**
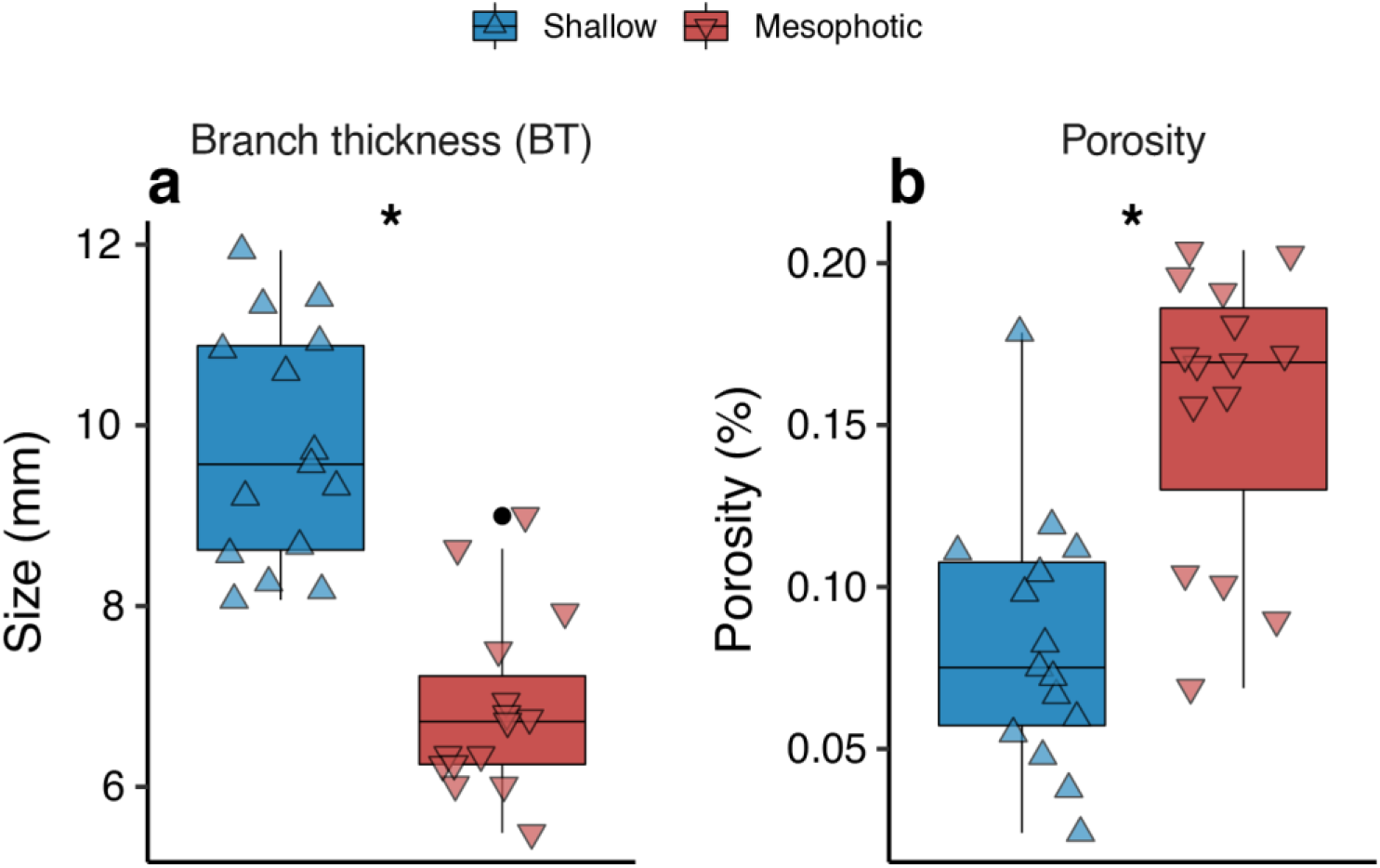
Box plots showing **(a)** branch thickness and **(b)** porosity between shallow (*blue*; triangle point up) and mesophotic (*red*; triangle point down) *Stylophora* corals (n = 15 per depth). Horizontal lines depict the median, box height depicts the interquartile range, whiskers depict ±1.5x interquartile range, and dots represent outliers. Asterisk denotes significance (p < 0.01).

**Figure 4.**
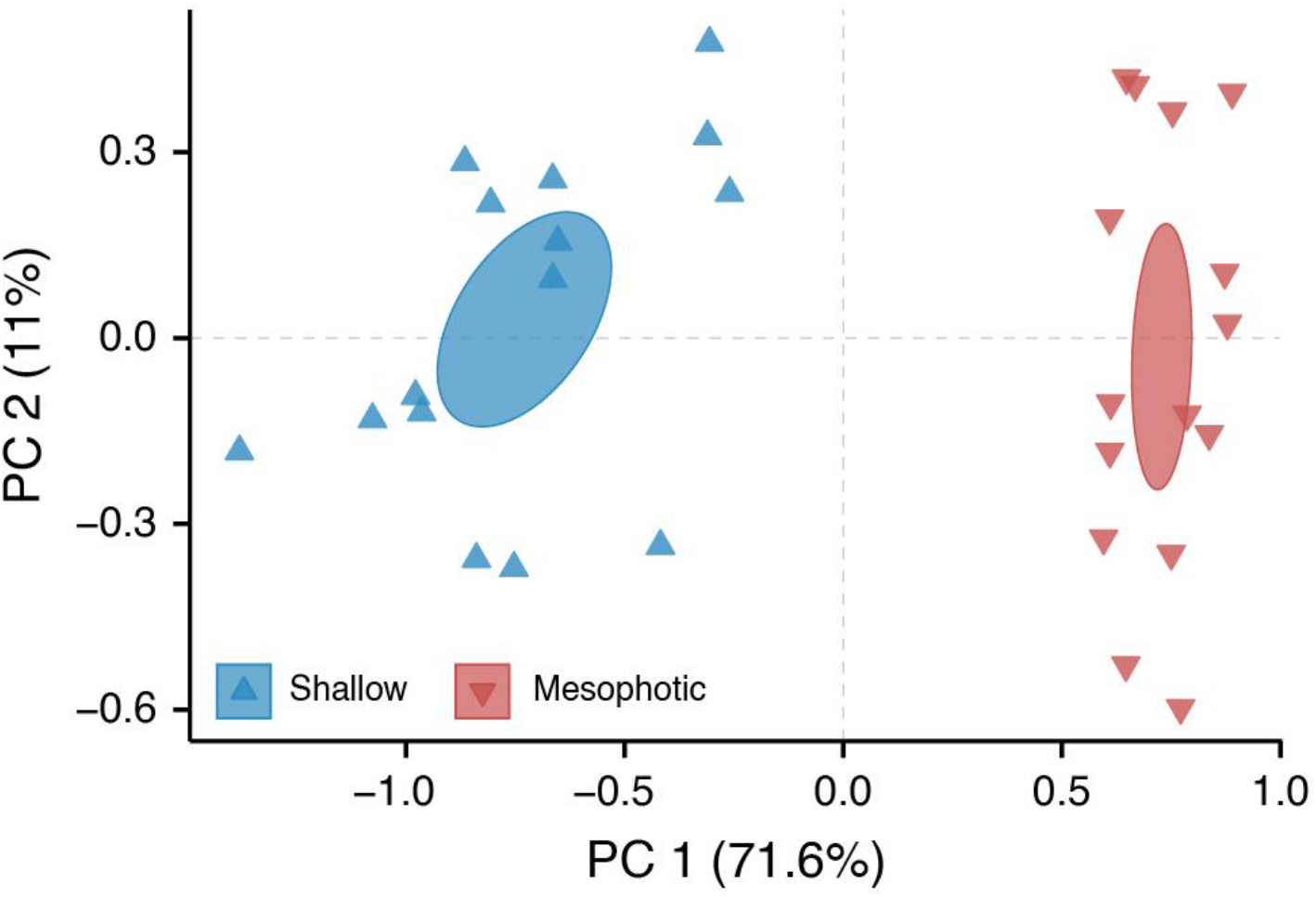
Principal coordinates analysis (PCoA) of the morphological characters of *S. pistillata* based on Euclidean space. Each color and shape represents a particular colony at a given depth (*n* = 30). Ellipses represent standard error.

Branch thickness was ∼30% thinner in mesophotic colonies than in shallow ones (MEPA, *p* < 0.01; Fig. 3a). Lastly, porosity analyses of mesophotic specimens revealed a 7.3% more porous skeleton than in shallow specimens, presenting 8.28 ± 0.01% and 15.57 ± 0.01% (mean ± SE), respectively (MEPA, *p* < 0.01; Fig. 3b).

### 3D models of light capture and photosynthesis

Based on the results of the morphometric analyses and the optical data from Kramer et al. (2021), we performed a total of 96 optical simulations (Figs. 5-6). Generally, normalized photosynthetic scores (*P*) under high-light simulations exhibited a wider range of values (*P* = 0.72-16.27) and displayed greater differences between shallow and mesophotic morphotypes than under low-light simulations (*P* = 5.64-13.94). The photosynthetic scores of shallow morphologies were dominated by an exponential decrease in fluence rate, while light attenuation was more homogenous for mesophotic corals (Fig. 5). Regardless of the *P-E* performance input (shallow and mesophotic), in nearly all simulation scenarios under low-light (45 µmol photons m^-2^ s^-1^) the photosynthetic scores of mesophotic morphotypes exceeded those of their shallow counterparts by up to 30% (Fig. 6a). In contrast, differences between morphotypes under all high-light scenarios (750 µmol photons m^-2^ s^-1^) were an order of magnitude higher in shallow versus mesophotic *P-E* performance inputs (*P* = 6.96-16.27 and 0.72-8.56, respectively; Fig. 6b). In most of the high-light simulation scenarios, shallow morphotypes exhibited 16-26% higher score values compared to the mesophotic morphotypes. For example, in the low µ_a_ tissue with shallow *P-E* parameters, photosynthetic scores were 15% higher for mesophotic morphotypes under low-light, while under high-light the score was 40% higher for the shallow morphotypes.

**Figure 5.**
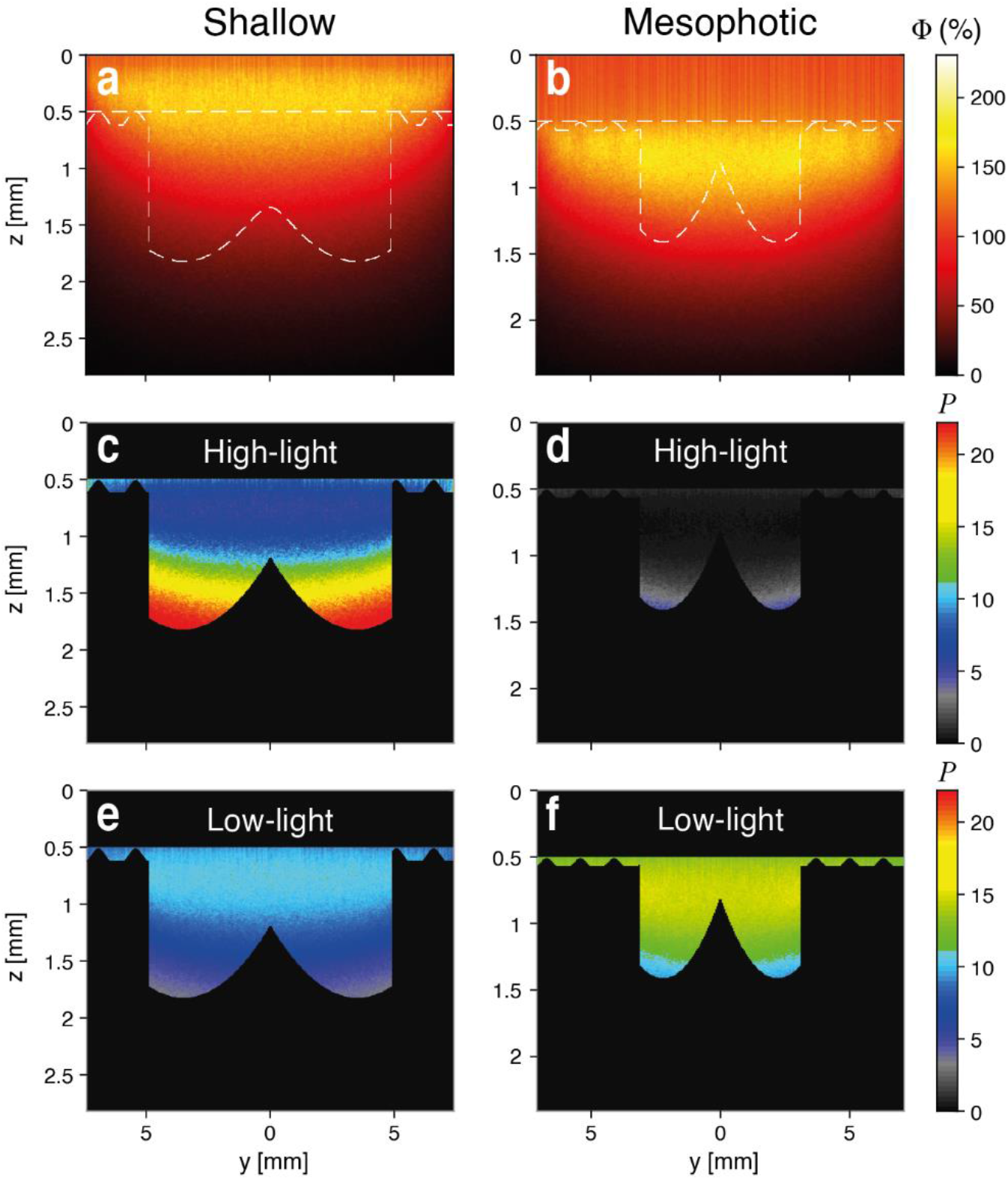
Examples of light propagation simulations shown in 2D (y-z axes) for simplicity (see normalized scores for all 96 scenarios in Fig. 5). **(a, b)** Relative fluence rates (Φ; delivered as W m^-2^; as color gradient) with contour indicating the surface boundaries of shallow and mesophotic natural morphotypes. **(c-f)** Photosynthetic score (*P*; as color gradient) on the tissue layer of shallow and mesophotic architectures with default settings under light intensities of 750 (high-light) and 45 (low-light) µmol photons m^-2^ s^-1^.

**Figure 6.**
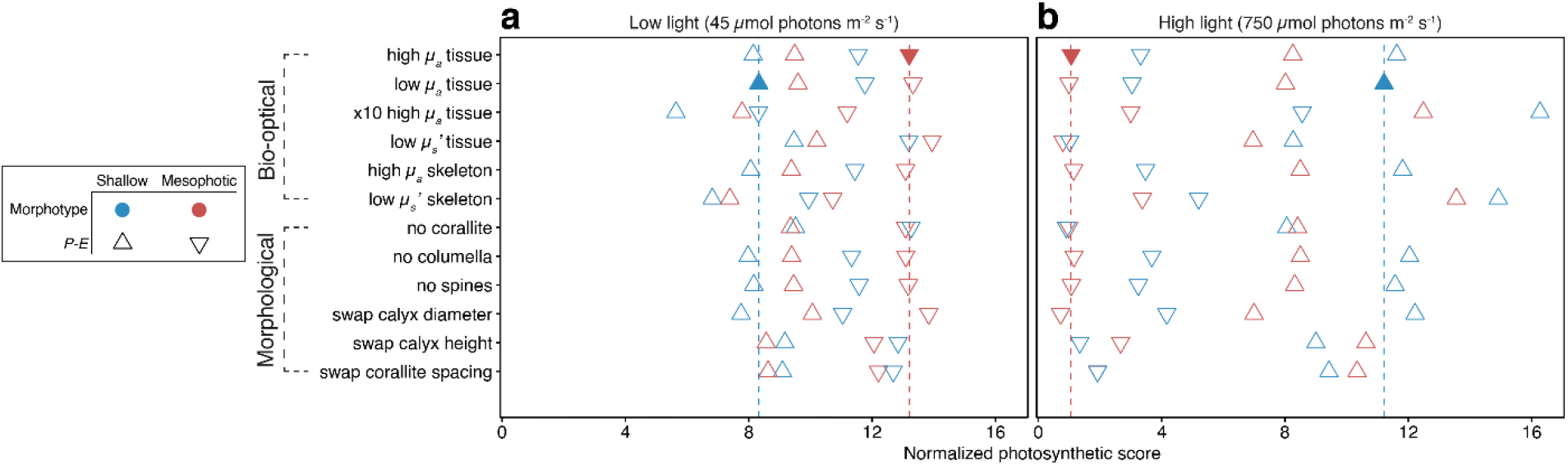
Normalized photosynthetic scores of different bio-optical and morphological simulation scenarios under **(a)** low-light (equivalent to 50 m; 45 µmol photons m^-2^ s^-1^) and **(b)** high-light (equivalent to 5m; 750 µmol photons m^-2^ s^-1^) conditions. Color denotes the morphotype and shape represents the ambient *P-E* performance. Filled triangles and dashed vertical lines represent the scores for default settings (high *µ*_*a*_ tissue, high *µ*_*s*_*’* tissue, low *µ*_*a*_ skeleton, high *µ*_*s*_*’* skeleton).

In contrast to the patterns noted above, exchanging calyx height and corallite spacing values between shallow and mesophotic morphotypes moderately increased the photosynthetic scores for shallow morphotypes under low-light; whereas under high-light conditions there was no difference between morphotypes exhibiting the mesophotic *P-E* parameters. Removing the corallite resulted in similar photosynthetic scores for both shallow and mesophotic morphotypes under both light conditions. Additionally, in most scenarios, surface complexity was greater in shallow morphologies, which exhibited an up to two-fold higher complexity than their mesophotic congeners (Table S2). However, exchanging corallite spacing or height between the two morphotypes resulted in similar surface complexities, which were akin to the mean value between the default morphotypes (Table S2).

## Discussion

Delineating the factors and functional traits that influence light capture by corals is fundamental for defining the range of light conditions under which survival, growth, and reproduction of a given coral species are possible. Using a mechanistic approach, we were able to uncover the role of key skeletal features of the coral *S. pistillata* in optimizing light harvesting. Our findings revealed morphology-based modifications adapted to local light conditions (i.e., shallow versus mesophotic), enabling optimized photon acquisition.

The multivariate analysis pertaining to the small-scale morphological traits revealed distinct morphotypes between corals of shallow and mesophotic origins (Fig. 4). Shallow corals were the most morphologically diverse group, while their mesophotic counterparts exhibited a narrower morphology diversity. Three dominant traits were shown to drive divergence along the first PCoA axis: calyx diameter, theca height, and corallite marginal spacing, which varied between depths in a coordinated way: the increase in corallite marginal spacing with depth had a strong negative correlation with the decrease in corallite size, while the corallite centers maintained their relative location in reference to their neighboring corallites (Fig. 2a, h, i). Notably, we demonstrate that in shallow-growing colonies the corallites expand in both width and depth and are closely spaced, while the opposite occurs in mesophotic corals (Fig. 2a, b, i). Additionally, we found that the coenosteum spines in mesophotic coral skeletons are significantly shorter and more closely spaced in comparison to those in the shallow depth (Fig. 2f). These findings are in line with earlier reports on the depth-related morphological changes in *S. pistillata* (Einbinder et al., 2009; Malik et al., 2020). Similar to our own findings, Ow and Todd (2010) reported that the calices of shallow *Goniastrea pectinata* fragments were deeper and the septae were shorter than in deeper fragments. However, these patterns are not consistent in all hermatypic coral species, since each species displays a distinct morphology with varying dimensions of the different skeletal features between deep and shallow depths. For example, in *Dipsastraea* (formerly *Favia speciosa*) and *Diploastrea heliopora*, the corallites expand and deepen, but are more spaced under shallow-water conditions (Todd et al., 2004); in *Galaxea facicularis*, corallite height increases and distance decreases with increasing light intensities, while corallite size increases under low-light levels (Crabbe and Smith, 2006); and in *Montastrea cavernosa*, the corallites are smaller and more spaced in mesophotic corals, while septal length decreases in their shallower counterparts (Studivan et al., 2019). Taken together with our current findings, these reports indicate that variation in small-scale skeletal geometry across light regimes is species specific. Consequently, it is unsound to draw generalized conclusions regarding shared skeletal features across coral species.

The coral host scatters light within its tissue by means of specialized tissue and skeletal modifications (Enríquez et al., 2005; Wangpraseurt et al., 2012), which increases the probability of photon absorption by the coral’s symbiotic microalgae (Wangpraseurt et al., 2016). Previous studies have demonstrated the effectiveness of two-dimensional models for investigating the interaction between light and coral architecture on a colony scale (Anthony et al., 2005; Muko et al., 2000) and on a single corallite scale (Ow and Todd, 2010). However, understanding how the different mechanisms of photoadaptation (e.g., morphological, physiological, and optical) interact to influence photosynthesis under a specific light regime is critical in determining the photic boundaries of any particular coral species. Integrating our morphometric results with recently obtained photosynthetic and optical data (Kramer et al., 2021), and using three-dimensional light propagation models, we applied a novel method by which to determine the functional significance of small-scale morphological traits with respect to the coral’s internal irradiance distribution.

Our simulation results demonstrate that small-scale morphological traits control *in-hospite* light absorption and coral photosynthetic performance. The change in the length-scale of morphological traits found within each of the two depth groups was shown to benefit the photosynthetic score with respect to their natural surrounding light regime (Figs. 5, 6). Overall, samples from shallow depths exhibited a more rapid attenuation of light and a greater ability to cope with excess light under high intensities, given that above the tissue surface the escaping flux (Φ) was enhanced by an up to two-fold higher incident irradiance (Fig. 5), thus supporting previous ecophysiological observations of light-adapted photosynthetic performance (Kramer et al., 2021; Martinez et al., 2020). On the colony scale, Hoogenboom et al. (2008) found evidence of a strong reduction in energy available for coral growth under high-light levels, and suggested that corals avoid the costs of excessive light exposure by means of altering colony morphology. Similarly, we show that the increase in corallite depth with increasing light intensities results in greater self-shading, thus providing an effective mechanism for keeping irradiance within a photophysiologically optimal range (Fig. S1). In contrast, the mesophotic architecture exhibited a more spacious structure, with a surface complexity reduced by nearly two-fold, which was advantageous in capturing low light. Hence, the combination of smaller, shallower, and more spaced corallites allowed for more light to be captured and utilized for photosynthesis (Fig. 2, 5). This principle appears to be valid for differential light gradients within the colony itself, as recently shown by Drake et al. (2021): corallites exposed to more light (i.e., at the tip of the branch) were less spaced and larger than corallites at the base and junction of the branch. The greater space occupied by the coenosteum relative to the corallites, as documented for mesophotic-depth colonies, may reflect the host’s response to minimize light limitation for its photosymbionts. This response may reduce the denser pigmentation of the polyps, as the polyps reveal the largest pigmentation cross-section when all the tentacles are retracted (Kramer et al., 2021; Wangpraseurt et al., 2012).

Surprisingly, simulations removing the corallites from the surface architecture yielded similar photosynthetic scores for the two morphotypes under the two light conditions (Fig. 6). Furthermore, surface complexity was found to be similar for the two morphotypes when exchanging corallite spacing and height values (Table S2). This exchange moderately increased the photosynthetic outcome, i.e., promoting a better photosynthetic advantage for shallow morphotypes under mesophotic light conditions, while the opposite occurred under shallow-water irradiance for mesophotic morphotypes. Several studies have described the important implications of coral structural complexity for light distribution. In large-scale structures, variation in colony surface complexity is related to competition and resource use, in which colonies whose surface distribution is complex have less light per unit surface area (Zawada et al., 2019). Similarly, a higher complexity in small-scale structures increases self-shading (Klaus et al., 2007; Ow and Todd, 2010; Wangpraseurt et al., 2012), as demonstrated in the shallow morphotypes of the present study. Consequently, we suggest that the corallite constitutes a dominant structural component, influencing surface complexity and subsequently light harvesting. In contrast, skeletal features such as the columella and coenosteal spines were shown to have a negligible impact on photosynthesis, indicating that their main role may be to provide additional structural and mechanical support to the coral tissue.

Typically, in comparison to shallow depths, depth-generalist corals in MCEs exhibit reduced growth rates (Groves et al., 2018; Mass et al., 2007) and lower reproductive performances (Shlesinger et al., 2018), assumingly due to light being a strong limiting energy source. Our findings highlight the fact that without specialized morphological modifications, light levels in MCEs would be insufficient to support the levels of photosynthesis required to sustain coral growth and reproduction (Fig. 6). Too much light would lead to photoinhibition; while too little light would not be sufficient to supply the corals’ nutrient demands. In terms of physiological adaptation, light-adapted photosymbionts exhibit well-developed photo-protective mechanisms, such as high NPQ levels (i.e., higher excess energy dissipation) and increased antioxidant capacity, while the symbiotic microalgae residing in mesophotic corals exhibit a highly efficient photosynthetic functioning (Einbinder et al., 2016; Martinez et al., 2020). However, light-driven physiological changes often occur in parallel with changes in the host characteristics, since the light field of Symbiodiniaceae populations is highly dependent on tissue thickness, corallite complexity, and optical properties (Enríquez et al., 2017; Kaniewska et al., 2011; Wangpraseurt et al., 2014). A recent study, for example, found that light-amplifying mechanisms in the host’s skeleton complement the photosynthetic demands of the photosymbionts (Kramer et al., 2021). In corals, light amplification is modulated by varying the scale of skeleton-length structures, ranging from nanometers (e.g., CaCO3 nanograins) to millimeters (e.g., corallite) (Swain et al., 2018). Hence, the skeleton geometry plays a vital role in dissipating adequate light to the tissue, as it controls the amount of energy that corals have available for growth and reproduction. Hoogenboom et al. (2008) posited that at the boundaries of the depth distribution, photoacclimation (i.e., physiological plasticity) cannot compensate for changes in morphology, and an adjustment of colony skeletal form appears to be the dominant phenotypic response; whereas photoacclimation is more important at intermediate depths. In line with that study, we suggest that the impact of symbiont physiology on mesophotic coral light acclimation is lower compared to its greater impacts on small-scale host morphology, as evidenced by our light simulation results.

In addition to phenotypic plasticity, morphological variability can also result from genetic influences (Bongaerts and Smith, 2019). To date, only a few studies have examined depth-related genetic partitioning in coral populations, demonstrating distinct patterns of vertical connectivity among species (Bongaerts et al., 2017, 2010; Serrano et al., 2016). Our study species, *S. pistillata*, was previously assessed for genetic vertical connectivity and found to belong to the same clade throughout its depth gradient in the Red Sea (Malik et al., 2020). The existing skeletal variations between depths are therefore reflective of genetic connectivity. As such, the smaller skeletal proportions in mesophotic corals may be a result of energy efficiency favoring reduced investment in skeletal features, arguably due to lower calcification rates (Anthony and Hoegh-Guldberg, 2003; Mass et al., 2007). Notwithstanding these energetic restraints, minimal energetic use is required to form the smaller mesophotic structures compared to the well-developed shallow architecture, since the need to create self-shading microhabitats is minimized in low-light environments.

Our results indicate that mesophotic *S. pistillata* skeletons exhibit a significantly greater porosity in comparison to their shallow congeners (Fig. 3b; see Fig. 1b,c vs e,f). Corals growing under decreased pH levels usually exhibit increased porosity due to reduced calcification rates (Fantazzini et al., 2015; Mollica et al., 2018). Similarly, the lower calcification rates of mesophotic *S. pistillata* colonies (Mass et al., 2007) may explain their increased porosity. A recent study by Fordyce et al. (2021), examined whether the endolithic microbial communities in coral skeletons may benefit from higher colony porosity since this potentially makes more space available for colonization in skeletal pores. However, they conclude that light capture by endoliths is affected by the material properties of the skeleton (i.e., density) and not by its porosity. We have shown here that the internal skeleton of mesophotic *S. pistillata* is more porous than its external engulfing-skeleton, and that the latter is thicker than the external skeleton of shallow-water branches (Fig. 1b,c,e,f). Given the imperforate nature of *S. pistillata* (i.e., its tissue does not penetrate the skeleton), we suggest that porosity in *S. pistillata* may be negligible in regard to light acquisition capability. However, unlike *S. pistillata*, the porous skeleton of perforate-tissue species may have a more significant function in light capture due to their tissues intercalating through the skeletal framework. Thus, we encourage future research into this issue in other coral species.

Although light energy is the primary energy source in the shallow waters (Muscatine, 1990), corals do not rely entirely on this form of energy. As mixotrophs, corals can acquire energy from multiple nutritional sources: namely, autotrophy – photosynthesis by photosymbionts; and heterotrophy – consuming zooplankton and particulate organic matter (Houlbrèque and Ferrier-Pagès, 2009). In shallow-water corals, heterotrophy can support survival during thermal stress by supplying energy to sustain symbiont autotrophy (Tremblay et al., 2016), while in some mesophotic species, heterotrophy can provide the host with an alternate source of energy in the lack of light (Lesser et al., 2010). However, since corallites of mesophotic *S. pistillata* colonies are significantly smaller than in their shallow congeners (Fig. 2a), this could potentially limit the size range of zooplankton available for capture. Nevertheless, Martinez et al. (2020) have shown that the photosynthesis pathway is the main source of carbon in both shallow and mesophotic *S. pistillata*, while heterotrophy represents a lower but similar portion of the total energy budget for both depths. Since quantitative changes in energy sources along the depth gradient are only known for a limited number of depth-generalists, with the findings being species-specific (Kahng et al., 2019), the role of heterotrophy as an energetic strategy at mesophotic depths remains to be further explored.

In conclusion, we have expanded the existing framework of light-harvesting strategies that allow corals to inhabit a wide range of light regimes. Specifically, our findings provide fundamental insights into how small-scale skeletal designs and properties modulate photosynthesis. The consensus in the literature is that changes in whole colony structure compensate for changes in light intensity along depth gradients (reviewed in Todd, 2008). In accord with those studies, we present evidence of morphology-based photoplasticity in *S. pistillata*, enabling an optimized small-scale skeletal adaptive response to the amount of available light. Our 3D simulations have shown that regardless of the optical modifications, mesophotic coral morphological traits consistently promoted a more effective light acquisition for photosynthesis under low-light simulations; while shallow coral morphological traits were better structured to cope with the high-light intensities they encounter. These findings indicate that small-scale morphological modifications constitute a more essential component of photoacclimation than optical ones at the photic boundaries. Moreover, coral populations living on the threshold of their optimal environment and adapted to extreme conditions have become useful models by which to predict the future functioning of coral reefs in light of climate change. Our 3D light models, integrating morphological and optical traits, could thus be applied to improve predictive models of coral responses to environmental changes. Furthermore, our findings provide a basis for future developments of coral-inspired technologies (e.g., bio-photoreactors) for clean and renewable energy, so vital in reducing atmospheric greenhouse gases.

## Materials and Methods

### Coral sampling and preparation

The study was conducted at the coral reefs of the northern GoE/A, Red Sea. The scleractinian coral *Stylophora pistillata* was chosen as a model species for this study due to its importance as an eco-engineering species in the GoE/A. *S. pistillata* is a branching colony characterized by very small-immersed corallites arranged in a plocoid morph, exhibiting a solid style-like columella with six poorly developed septa, and a spiny coenosteum (Veron et al., 2000). Furthermore, it exhibits a wide bathymetric distribution (Kramer et al., 2020; Loya, 1976) and pronounced morphological variation in colony growth form with depth, from a subspherical densely-branched form in the shallows to a more spread-out branch morphology in mesophotic environments (Fig. 1).

Fragments from intact adult coral colonies (ca. 20-25 cm in diameter) were collected during recreational and closed-circuit rebreather dives from shallow (5 m) and upper mesophotic (45-50 m) depths, corresponding to 40-45% and 3-8% of midday surface PAR, respectively (Tamir et al., 2019). In total, fragments from 30 colonies were used for this study (*n* = 15 per depth). Conspecific coral colonies were sampled at least five meters apart to avoid sampling clone mates. The samples were submerged in 5% sodium hypochlorite (NaClO) for 24 hours to dissolve the soft tissue, rinsed with distilled water, and air-dried at room temperature.

### X-ray microtomography and morphometrics

For analysis of the morphometric characters, each sample was scanned using high-resolution micro-computed tomography (μCT), conducted with a Nikon XT H 225ST µCT (Nikon Metrology Inc., USA) at The Steinhart Museum of Natural History, Tel Aviv University. *S. pistillata* specimens were scanned at an isotropic voxel (volume pixels) size of 10 μm (360° rotation), with voltage and current set to 170 kV and 56 µA, respectively. Scans from each specimen were saved in a TIFF image format for 3D volume rendering and quantitative analysis using the software Dragonfly (© 2021 Object Research System (ORS) Inc.).

All measurements were taken from random intact corallites and from the coenosteum surrounding them, and which were not in a budding state nor at the colony margins (at least 2 cm from the distal branch tip to avoid areas of recent growth). A total of ten small-scale (mm) skeletal morphometric traits were measured (≥ 10 measurements per trait per sample; Fig. 1): calyx diameter (CD), theca (corallite wall) height (TH), septal length (SL), septa width (SW), columella height (CH), coenosteum spinule spacing (SS), coenosteum spinule length (SPL), coenosteum spinule width (SPW), and coral spacing, which was measured in two ways: the distance between neighboring corallite centers (CSC) and minimal distance between neighboring corallites (CSM). An additional measurement comprised branch thickness (mm). All skeletal metrics were perpendicularly aligned to the sample’s growth axis prior to measurement. Lastly, apparent porosity was determined as the percentage ratio of pore volume to the total volume occupied by the coral skeleton.

### 3D light propagation models

To model the effect of different skeletal features on light capture we developed a 3D Monte Carlo simulation (Jacques et al., 2013; Wangpraseurt et al., 2016). Monte Carlo Simulations are probability distribution models that are widely used for modeling light propagation in biological tissues and often considered the gold standard for modeling complex tissue architectures (Tuchin, 2015). Detailed explanations of the core simulation process can be found in Wang et al. (1995). Briefly, photons are launched through a tissue with independent absorption and scattering centers, and interact with the tissue via a random process of light scattering and absorption. The overall probability of absorption and scattering are based on the inherent optical properties of the tissues of interest, yielding a characteristic, average light distribution. Monte Carlo Simulations allow for modeling any source geometry, with mesh-based and voxel-based methods existing for modeling complex 3D geometries.

### Source architecture

We used the average morphological parameters obtained from µCT scanning to create representative coral skeleton designs for shallow and mesophotic corals (Fig. 1, Table S1). For the coral tissue, we assumed thicknesses based on previous measurements (Kramer et al., 2021). It is important to note that coral tissues are flexible, and expansion and contraction can affect light propagation. For simplicity, we assumed here only the contracted tissue state, comprising one continuous tissue type with average optical properties (see below). The tissue covering the coenosteum was set to the maximal length of the coenosteal spines, and filled the calyx cavity to mimic a fully contracted coral polyp. The void space was filled with water.

### Simulation settings

We conducted a series of simulation scenarios to assess the importance of different morphological and optical traits for coral light capture. First, we conducted two simulations for each morphotype with identical optical properties (see Table S3 for an overview). We then assessed the contribution of individual architectural features using a “knock-out” procedure, which involves removing one morphological trait at a time and assessing the light distribution over the entire coral architecture. For the simulation in which the calyx was removed, we kept the tissue volume constant by redistributing the tissue over the coenosteum. We further quantified the effects of morphological traits on surface area (mm^2^), surface complexity (geometric surface area divided by real surface area), and tissue volume (mm^3^) for shallow and mesophotic architectures. Moreover, we also exchanged the mean measurement values of the above traits between shallow and mesophotic morphotypes to further test their functionality. Finally, we examined the contribution of the skeletal architecture given the same optical properties, with each simulation scenario focusing on one modified optical trait.

To determine the optimal simulation time, we executed multiple tests with the same setting and varying simulation times. We found that simulations over two hours yielded similar results to those of the two-hour simulations, and thus decided to use two-hour simulations in all scenarios. With this setup, we executed the MC simulation code (2 hours/ ∼5×10^7^ photons; resolution = 0.005 mm/pixel) and obtained the fluence rate information on the 3D coral models.

### 3D photosynthesis model

To evaluate the relationship between coral light capture and coral photosynthesis we developed a novel 3D photosynthesis model. The model uses the volumetric fluence rate distribution to calculate tissue photosynthesis for complex coral architectures at a high spatial resolution. We developed a script to calculate a ‘relative photosynthesis score’ using our experimentally determined photosynthesis-irradiance (*P-E*) data from Pulse Amplitude Modulation (PAM) chlorophyll-*a* fluorometer (Fig. S1), and the following relationship (Ritchie and Larkum, 2012)(Table S1):

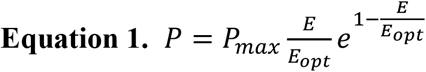

where *P* represents the relative gross photosynthesis score, *E* is the fluence rate, *P*_*max*_ represents the maximum gross photosynthesis rate, and *E*_*opt*_ is the optimal fluence rate at *P*_*max*_. Score values were normalized by tissue voxels for each morphotype. For each experimental setting, we calculated the actual fluence rates based on the *in-situ* light levels (Tamir et al., 2019): 45 and 750 µmol photons m^-2^ s^-1^ for shallow (5 m) and mesophotic (50 m) depths, respectively.

### Statistical analyses

Statistical analyses were performed using the R software (R Core Team 2021). Since in most cases the data did not conform to parametric test assumptions, intraspecies variations between depths for each morphological character were tested using a mixed-effects permutational analysis (MEPA; 999 permutations) and included the sample ID as a random effect. These analyses were run using the {lme4}(Bates et al., 2015) and {predictmeans}(Luo et al., 2021) packages. A principal coordinates analysis (PCoA) based on a Euclidean distance matrix of standardized data was created with the {vegan} package to visualize the pattern of morphological variation between depths in a multivariate trait space. Permutational multivariate analysis of variance (PERMANOVA; 999 permutations) was performed to determine the overall effect of depth on the morphological patterns. Traits were highlighted as important for a given axis based on whether their loadings exceeded the null contribution value of 10% (100% divided by ten variables). Pearson’s correlation coefficients were used to assess pairwise correlations among the different skeletal traits. Similarity percentage analysis (SIMPER) was conducted to determine which morphological traits were responsible for most of the variation between depths (Clarke, 1993).

## Acknowledgments

We are grateful to the Interuniversity Institute for Marine Sciences in Eilat (IUI) for making its facilities available to us. We thank S. Ellenbogen from the Dan David Center for Human Evolution and Biohistory Research for her technical assistance; K. Mumm and M. Bocanegra for their assistance with the light models; and N. Paz for editing the manuscript. This work was supported by the Israel Science Foundation (ISF) Grant No. 1191/16 to YL, and by the ASSEMBLE-Plus consortium grant to D.W.

## Author contributions

N.K., D.W., and Y.L. conceived and designed the research. N.K. collected the coral samples, conducted μCT work, performed data analysis, and wrote the first draft of the manuscript. N.K. and J.G. generated the figures. J.G. and D.W. developed the optical simulations. All authors contributed to editing the manuscript and gave final approval for publication.

## References

Anthony KRN, Hoegh-Guldberg O. 2003. Variation in coral photosynthesis, respiration and growth characteristics in contrasting light microhabitats: an analogue to plants in forest gaps and understoreys? Funct Ecol 17:246–259. doi:10.1046/j.1365-2435.2003.00731.x

Anthony KRN, Hoogenboom MO, Connolly SR. 2005. Adaptive variation in coral geometry and the optimization of internal colony light climates. Funct Ecol 19:17–26. doi:10.1111/j.0269-8463.2005.00925.x

Barnes D. 1973. Growth in colonial scleractinians. Bull Mar Sci 280–298.

Bates D, Mächler M, Bolker B, Walker S. 2015. Fitting Linear Mixed-Effects Models Using {lme4}. J Stat Softw 67:1–48. doi:10.18637/jss.v067.i01

Bongaerts P, Ridgway T, Sampayo EM, Hoegh-Guldberg O. 2010. Assessing the “deep reef refugia” hypothesis: Focus on Caribbean reefs. Coral Reefs 29:1–19. doi:10.1007/s00338-009-0581-x

Bongaerts P, Riginos C, Brunner R, Englebert N, Smith SR. 2017. Deep reefs are not universal refuges : reseeding potential varies among coral species. Sci Adv 3:1–40.

Bongaerts P, Smith TB. 2019. Beyond the “Deep Reef Refuge” Hypothesis: A Conceptual Framework to Characterize Persistence at Depth In: Loya Y, Puglise KA, Bridge TCL, editors. Mesophotic Coral Ecosystems. Cham: Springer International Publishing. pp. 881–895. doi:10.1007/978-3-319-92735-0_45

Bragg JG, Westoby M. 2002. Leaf size and foraging for light in a sclerophyll woodland. Funct Ecol 16:633–639. doi:https://doi.org/10.1046/j.1365-2435.2002.00661.x

Bruno JF, Edmunds PJ. 1997. Clonal Variation for Phenotypic Plasticity in the coral Madracis Mirabilis. Ecology 78:2177–2190. doi:10.1890/0012-9658(1997)078[2177:CVFPPI]2.0.CO;2

Chappell J. 1980. Coral morphology, diversity and reef growth. Nature 286:249–252. doi:10.1038/286249a0

Clarke KR. 1993. Non-parametric multivariate analyses of changes in community structure. Aust J Ecol 18:117–143. doi:10.1111/j.1442-9993.1993.tb00438.x

Crabbe MJC, Smith DJ. 2006. Modelling variations in corallite morphology of Galaxea fascicularis coral colonies with depth and light on coastal fringing reefs in the Wakatobi Marine National Park (S.E. Sulawesi, Indonesia). Comput Biol Chem 30:155–159. doi:https://doi.org/10.1016/j.compbiolchem.2005.11.004

Doszpot N, McWilliam M, Pratchett M, Hoey A, Figueira W. 2019. Plasticity in Three-Dimensional Geometry of Branching Corals Along a Cross-Shelf Gradient. Diversity 11:44. doi:10.3390/d11030044

Drake JL, Malik A, Popovits Y, Yosef O, Shemesh E, Stolarski J, Tchernov D, Sher D, Mass T. 2021. Physiological and Transcriptomic Variability Indicative of Differences in Key Functions Within a Single Coral Colony. Front Mar Sci 8:768. doi:10.3389/fmars.2021.685876

Drake JL, Mass T, Stolarski J, Von Euw S, van de Schootbrugge B, Falkowski PG. 2020. How corals made rocks through the ages. Glob Chang Biol. doi:10.1111/gcb.14912

Dubé CE, Mercière A, Vermeij MJA, Planes S. 2017. Population structure of the hydrocoral Millepora platyphylla in habitats experiencing different flow regimes in Moorea, French polynesia. PLoS One 12:1–20. doi:10.1371/journal.pone.0173513

Dubinsky Z, Falkowski PG, Porter JW, Muscatine L. 1984. Absorption and Utilization of Radiant Energy by Light-and Shade-Adapted Colonies of the Hermatypic Coral Stylophora pistillata. Proc R Soc B Biol Sci 222:203–214. doi:10.1098/rspb.1984.0059

Dustan P. 1982. Depth-dependent photoadaption by zooxanthellae of the reef coral Montastrea annularis. Mar Biol 68:253–264. doi:10.1007/BF00409592

Einbinder S, Gruber DF, Salomon E, Liran O, Keren N, Tchernov D. 2016. Novel Adaptive Photosynthetic Characteristics of Mesophotic Symbiotic Microalgae within the Reef-Building Coral, Stylophora pistillata. Front Mar Sci 3:195. doi:10.3389/fmars.2016.00195

Einbinder S, Mass T, Brokovich E, Dubinsky Z, Erez J, Tchernov D. 2009. Changes in morphology and diet of the coral Stylophora pistillata along a depth gradient. Mar Ecol Prog Ser 381:167–174. doi:10.3354/meps07908

Enríquez S, Méndez ER, Hoegh-Guldberg O, Iglesias-Prieto R. 2017. Key functional role of the optical properties of coral skeletons in coral ecology and evolution. Proc R Soc B Biol Sci 284. doi:10.1098/rspb.2016.1667

Enríquez S, Méndez ER, Iglesias-Prieto R. 2005. Multiple scattering on coral skeletons enhances light absorption by symbiotic algae. Limnol Oceanogr 50:1025–1032. doi:10.4319/lo.2005.50.4.1025

Falkowski PG, Dubinsky Z. 1981. Light-shade adaptation of Stylophora pistillata, a hermatypic coral from the Gulf of Eilat. Nature 289:172–174. doi:10.1038/289172a0

Fantazzini P, Mengoli S, Pasquini L, Bortolotti V, Brizi L, Mariani M, Di Giosia M, Fermani S, Capaccioni B, Caroselli E, Prada F, Zaccanti F, Levy O, Dubinsky Z, Kaandorp JA, Konglerd P, Hammel JU, Dauphin Y, Cuif JP, Weaver JC, Fabricius KE, Wagermaier W, Fratzl P, Falini G, Goffredo S. 2015. Gains and losses of coral skeletal porosity changes with ocean acidification acclimation. Nat Commun 6. doi:10.1038/ncomms8785

Graham NAJ, Nash KL. 2013. The importance of structural complexity in coral reef ecosystems. Coral Reefs. doi:10.1007/s00338-012-0984-y

Grottoli AG, Warner ME, Levas SJ, Aschaffenburg MD, Schoepf V, Mcginley M, Baumann J, Matsui Y. 2014. The cumulative impact of annual coral bleaching can turn some coral species winners into losers. Glob Chang Biol 20:3823–3833. doi:10.1111/gcb.12658

Groves SH, Holstein DM, Enochs IC, Kolodzeij G, Manzello DP, Brandt ME, Smith TB. 2018. Growth rates of Porites astreoides and Orbicella franksi in mesophotic habitats surrounding St. Thomas, US Virgin Islands. Coral Reefs 37:345–354. doi:10.1007/s00338-018-1660-7

Hoogenboom MO, Connolly SR, Anthony KRN. 2008. Interactions between morphological and physiological plasticity optimize energy acquisition in corals. Ecology 89:1144–1154. doi:https://doi.org/10.1890/07-1272.1

Houlbrèque F, Ferrier-Pagès C. 2009. Heterotrophy in tropical scleractinian corals. Biol Rev 84:1–17. doi:10.1111/j.1469-185X.2008.00058.x

House JE, Brambilla V, Bidaut LM, Christie AP, Pizarro O, Madin JS, Dornelas M. 2018. Moving to 3D: relationships between coral planar area, surface area and volume. PeerJ 6:e4280. doi:10.7717/peerj.4280

J. FA, D. AT W. L, Katherine M. 2021. Light Capture, Skeletal Morphology, and the Biomass of Corals’ Boring Endoliths. mSphere 6:e00060–21. doi:10.1128/mSphere.00060-21

Jacques S, Li T, Prahl S. 2013. mcxyz. c, a 3D Monte Carlo simulation of heterogeneous tissues. omlcorg/software/mc/mcxyz.

Kahng SE, Akkaynak D, Shlesinger T, Hochberg EJ, Wiedenmann J, Tamir R, Tchernov D. 2019. Light, Temperature, Photosynthesis, Heterotrophy, and the Lower Depth Limits of Mesophotic Coral Ecosystems In: Loya Y, Puglise KA, Bridge TCL, editors. Mesophotic Coral Ecosystems. Cham: Springer International Publishing. pp. 801–828. doi:10.1007/978-3-319-92735-0_42

Kahng SE, Copus JM, Wagner D. 2014. Recent advances in the ecology of mesophotic coral ecosystems (MCEs). Curr Opin Environ Sustain 7:72–81. doi:https://doi.org/10.1016/j.cosust.2013.11.019

Kahng SE, Watanabe TK, Hu H-M, Watanabe T, Shen C-C. 2020. Moderate zooxanthellate coral growth rates in the lower photic zone. Coral Reefs. doi:10.1007/s00338-020-01960-4

Kaniewska P, Magnusson SH, Anthony KRN, Reef R, Kühl M, Hoegh-Guldberg O. 2011. Importance of macro-versus microstructure in modulating light levels inside coral colonies. J Phycol 47:846–860. doi:https://doi.org/10.1111/j.1529-8817.2011.01021.x

Klaus JS, Budd AF, Fouke B. 2007. Environmental controls on corallite morphology in the reef coral Montastraea annularis Hot springs microbiology View project Positive Accretion in the Deep: Carbonate budget analysis of Caribbean mesophotic coral reef habitats View project. Bull Mar Sci 80:233–260.

Kramer N, Tamir R, Ben-Zvi O, Jacques SL, Loya Y, Wangpraseurt D. 2021. Light-harvesting in mesophotic corals is powered by a spatially efficient photosymbiotic system between coral host and microalgae. bioRxiv 2020.12.04.411496. doi:10.1101/2020.12.04.411496

Kramer N, Tamir R, Eyal G, Loya Y. 2020. Coral Morphology Portrays the Spatial Distribution and Population Size-Structure Along a 5–100 m Depth Gradient. Front Mar Sci 7:615. doi:10.3389/fmars.2020.00615

Laverick JH, Tamir R, Eyal G, Loya Y. 2020. A generalized light-driven model of community transitions along coral reef depth gradients. Glob Ecol Biogeogr 29:1554–1564. doi:10.1111/geb.13140

Lesser MP, Marc S, Michael S, Michiko O, Gates RD, Andrea G. 2010. Photoacclimatization by the coral Montastraea cavernosa in the mesophotic zone: Light, food, and genetics. Ecology. doi:10.1890/09-0313.1

Lichtenthaler HK, Ač A, Marek M V, Kalina J, Urban O. 2007. Differences in pigment composition, photosynthetic rates and chlorophyll fluorescence images of sun and shade leaves of four tree species. Plant Physiol Biochem 45:577–588. doi:https://doi.org/10.1016/j.plaphy.2007.04.006

Loya Y. 1976. The Red Sea coral Stylophora pistillata is an r strategist. Nature 259:478–480. doi:10.1038/260170a0

Luo D, Ganesh S, Koolaard J. 2021. predictmeans: Calculate Predicted Means for Linear Models.

Malik A, Einbinder S, Martinez S, Tchernov D, Haviv S, Almuly R, Zaslansky P, Polishchuk I, Pokroy B, Stolarski J, Mass T. 2020. Molecular and skeletal fingerprints of scleractinian coral biomineralization: From the sea surface to mesophotic depths. Acta Biomater 1–14. doi:10.1016/j.actbio.2020.01.010

Martinez S, Kolodny Y, Shemesh E, Scucchia F, Nevo R, Levin-Zaidman S, Paltiel Y, Keren N, Tchernov D, Mass T. 2020. Energy Sources of the Depth-Generalist Mixotrophic Coral Stylophora pistillata. Front Mar Sci 7:1–16. doi:10.3389/fmars.2020.566663

Mass T, Einbinder S, Brokovich E, Shashar N, Vago R, Erez J, Dubinsky Z. 2007. Photoacclimation of Stylophora pistillata to light extremes: Metabolism and calcification. Mar Ecol Prog Ser 334:93–102. doi:10.3354/meps334093

Mollica NR, Guo W, Cohen AL, Huang K-F, Foster GL, Donald HK, Solow AR. 2018. Ocean acidification affects coral growth by reducing skeletal density. Proc Natl Acad Sci 115:1754–1759. doi:10.1073/pnas.1712806115

Muko S, Kawasaki K, Sakai K, Takasu F, Shigesada N. 2000. Morphological plasticity in the coral Porites sillimaniani and its adaptive significance. Bull Mar Sci 66:225–239.

Munday P, Jones G. 1998. The ecological implications of small body size among coral-reef fishes. Oceanogr Mar Biol 36:373–411.

Muscatine L. 1990. The role of symbiotic algae in carbon and energy flux in coral reefs. Ecosyst world 25:75–87.

Ow YX, Todd PA. 2010. Light-induced morphological plasticity in the scleractinian coral Goniastrea pectinata and its functional significance. Coral Reefs 29:797–808. doi:10.1007/s00338-010-0631-4

Paz-García DA, Aldana-Moreno A, Cabral-Tena RA, García-De-León FJ, Hellberg ME, Balart EF. 2015. Morphological variation and different branch modularity across contrasting flow conditions in dominant Pocillopora reef-building corals. Oecologia 178:207–218. doi:10.1007/s00442-014-3199-9

Pereira PHC, Munday PL. 2016. Coral colony size and structure as determinants of habitat use and fitness of coral-dwelling fishes. Mar Ecol Prog Ser 553:163–172.

Ritchie RJ, Larkum AWD. 2012. Modelling photosynthesis in shallow algal production ponds. Photosynthetica 50:481–500. doi:10.1007/s11099-012-0076-9

Sæbø A, Krekling T, Appelgren M. 1995. Light quality affects photosynthesis and leaf anatomy of birch plantlets in vitro. Plant Cell Tissue Organ Cult 41:177–185. doi:10.1007/BF00051588

Serrano XM, Baums IB, Smith TB, Jones RJ, Shearer TL, Baker AC. 2016. Long distance dispersal and vertical gene flow in the Caribbean brooding coral Porites astreoides. Sci Rep 6:21619. doi:10.1038/srep21619

Sherman CE, Locker SD, Webster JM, Weinstein DK. 2019. Geology and Geomorphology In: Loya Y, Puglise KA, Bridge TCL, editors. Mesophotic Coral Ecosystems. Cham: Springer International Publishing. pp. 849–878. doi:10.1007/978-3-319-92735-0_44

Shlesinger T, Grinblat M, Rapuano H, Amit T, Loya Y. 2018. Can mesophotic reefs replenish shallow reefs? Reduced coral reproductive performance casts a doubt. Ecology. doi:10.1002/ecy.2098

Smith LW, Barshis D, Birkeland C. 2007. Phenotypic plasticity for skeletal growth, density and calcification of Porites lobata in response to habitat type. Coral Reefs 26:559–567. doi:10.1007/s00338-007-0216-z

Soto D, De Palmas S, Ho MJ, Denis V, Chen CA. 2018. Spatial variation in the morphological traits of Pocillopora verrucosa along a depth gradient in Taiwan. PLoS One 13:1–20. doi:10.1371/journal.pone.0202586

Studivan MS, Milstein G, Voss JD. 2019. Montastraea cavernosa corallite structure demonstrates distinct morphotypes across shallow and mesophotic depth zones in the Gulf of Mexico. PLoS One 14. doi:10.1371/journal.pone.0203732

Swain TD, Lax S, Lake N, Grooms H, Backman V, Marcelino LA. 2018. Relating Coral Skeletal Structures at Different Length Scales to Growth, Light Availability to Symbiodinium, and Thermal Bleaching. Front Mar Sci 5. doi:10.3389/fmars.2018.00450

Tamir R, Eyal G, Kramer N, Laverick JH, Loya Y. 2019. Light environment drives the shallow to mesophotic coral community transition. Ecosphere 10:e02839. doi:10.1101/622191

Team RC. 2021. R: A Language and Environment for Statistical Computing. R Foundation for Statistical Computing, Vienna, Austria. URL https://www.R-project.org/.

Todd PA. 2008. Morphological plasticity in scleractinian corals. Biol Rev. doi:10.1111/j.1469-185X.2008.00045.x

Todd PA, Ladle RJ, Lewin-Koh NJI, Chou LM. 2004. Genotype x environment interactions in transplanted clones of the massive corals Favia speciosa and Diploastrea heliopora. Mar Ecol Prog Ser. doi:10.3354/meps271167

Tremblay P, Gori A, Maguer JF, Hoogenboom M, Ferrier-Pagès C. 2016. Heterotrophy promotes the re-establishment of photosynthate translocation in a symbiotic coral after heat stress. Sci Rep 6:38112. doi:10.1038/srep38112

Tuchin V V. 2015. Tissue Optics: Light Scattering Methods and Instruments for Medical Diagnostics, Third Edit. ed. SPIE PRESS.

Veron C, Stafford-Smith M, Turak E, DeVantier L. 2000. Corals of the world.

Wang L, Jacques SL, Zheng L. 1995. MCML—Monte Carlo modeling of light transport in multi-layered tissues. Comput Methods Programs Biomed 47:131–146. doi:https://doi.org/10.1016/0169-2607(95)01640-F

Wangpraseurt D, Jacques SL, Petrie T, Kühl M. 2016. Monte Carlo Modeling of Photon Propagation Reveals Highly Scattering Coral Tissue. Front Plant Sci 7:1–10. doi:10.3389/fpls.2016.01404

Wangpraseurt D, Larkum AWD, Ralph PJ, Kühl M. 2012. Light gradients and optical microniches in coral tissues. Front Microbiol 3:1–9. doi:10.3389/fmicb.2012.00316

Wangpraseurt D, Polerecky L, Larkum AWD, Ralph PJ, Nielsen DA, Pernice M, Kühl M. 2014. The in situ light microenvironment of corals. Limnol Oceanogr 59:917–926. doi:10.4319/lo.2014.59.3.0917

Wehrberger F, Herler J. 2014. Microhabitat characteristics influence shape and size of coral-associated fishes. Mar Ecol Prog Ser 500:203–214. doi:10.3354/meps10689

Willis BL. 1985. Phenotypic plasticity versus phenotypic stability in the reef corals Turbinaria mesenterina and Pavona cactus. Proc Fifth Int Coral Reef Symp 4:107–112.

Zawada Kyle J. A., Dornelas M, Madin JS. 2019. Quantifying coral morphology. Coral Reefs 38:1281–1292. doi:10.1007/s00338-019-01842-4

Zawada Kyle J.A., Madin JS, Baird AH, Bridge TCL, Dornelas M. 2019. Morphological traits can track coral reef responses to the Anthropocene. Funct Ecol 33:962–975. doi:10.1111/1365-2435.13358

